# Frequent subgraph mining for biologically meaningful structural motifs

**DOI:** 10.1101/2020.05.14.095695

**Authors:** Sebastian Keller, Pauli Miettinen, Olga V. Kalinina

## Abstract

Identification of biologically relevant motifs in proteins is a long-standing problem in bioinformatics, especially when considering distantly related proteins where sequence analysis alone becomes increasingly difficult. Here we present a novel approach to identify such motifs in protein three-dimensional structures without depending on sequence alignment by representing structures as graphs in the form of residue interaction networks and employing a modified frequent subgraph mining algorithm. These networks represent residues as vertices while contacts between residues are denoted by edges labeled with Euclidean distances. We use frequent subgraph mining to determine all subgraphs that are subgraph isomorphic to, i.e. are contained in, at least a given number of such networks generated from structures in the same protein family. For this we introduce two extensions of the classical frequent subgraph mining: approximate matching of distance-based labels to account for small variations between protein structures and scoring as well as score-based filtering of subgraphs in order to identify structurally conserved motifs and to counteract the expanding size of the search space. This approach was then validated by demonstrating that it can rediscover previously characterized functionally important structural motifs in selected protein families. For further validation we show that it is also able to identify motifs that correspond to patterns in the PROSITE database. We then applied our approach to all superfamilies in the SCOP database and found an enrichment of residues in the ligand binding site in the discovered motifs evidencing their functional importance. Finally we use the approach to discover a novel structural motif in jelly-roll capsid proteins found in members of the picornavirus-like superfamily. This is presented together with an efficient open source implementation of the algorithm called RINminer.

**Author summary:** As the evolutionary distance between proteins increases, their sequence identity drops rapidly, whereas functionally important sequence motifs and three-dimensional (3D) structural scaffold, in which they are embedded, are more conserved. We developed an approach that automatically identifies such motifs by converting protein 3D structures into a set of graphs and then employing the frequent subgraph mining framework. In these graphs, residues are represented as vertices, and if two residues interact in the corresponding protein 3D structure, they are connected by an edge labeled with the Euclidean distance between the residues. In the classical setting of frequent subgraph mining, all subgraphs from a database of graphs are enumerated and the ones that are exactly found, i.e. are subgraph isomorphic, in more than a certain number of graphs are listed as supported. Our approach introduces two new concepts: approximately isomorphic subgraphs and an efficient scoring scheme that allows to retain only biologically relevant subgraph in the enumeration step. Approximate isomorphism allows edge labels not to match exactly, and thus account for natural deviations between 3D structures of related proteins. With our approach, we were able to automatically rediscover known motifs from PROSITE, as well as in three well-studied extremely diverse protein families. We predicted functionally important residues in SCOP superfamilies and demonstrated that they tend to lie in structurally meaningful regions: ligand-binding sites and protein core. Additionally, we present a previously unreported structural motif in jelly-roll viral capsids.

## 1 Introduction

The three-dimensional (3D) structure of distantly related proteins is often more conserved than their amino acid sequence [1]. Such proteins may belong to the same family, e.g. kinases, but from distantly related organisms. Also in rapidly evolving organisms, such as viruses, very diverse protein groups can be found that are difficult to approach with sequence analysis tools. Discovering functionally important motifs in sequences or structures of these protein families is a challenging task that often requires manual intervention and expert knowledge. One example of such motifs conserved in a family of proteins with very low sequence similarity is found in viral RNA-dependent RNA polymerases (RdRPs) [2]. Several RdRP sequence motifs, including those common to RdRPs from positive and negative single-strand RNA viruses as well as retroviruses, were identified before the availability of any 3D structures [3]. Later resolved 3D structures of these proteins confirmed that these sequence motifs that lie at variable distances in the RdRP sequences from different species all cluster together near or directly in the active site of the protein [2]. With 3D structures at hand, it now is more reasonable to consider this as a single structural motif rather than a set of sequence motifs with variable spacing between them.

In this study, we present a novel algorithm for the detection of such structural motifs based on 3D structures and validate it using large data sets of related proteins from the PROSITE [4] and SCOP [5] databases. Our method is based on a solid graph-theoretical and data-mining foundation, and has been implemented as an efficient tool in C called *RINminer*, available at https://github.com/kalininalab/rinminer. We demonstrate a good agreement of motifs identified with RINminer with functional patterns in PROSITE. For SCOP superfamilies, we show a significant enrichment of residues from the identified motifs in functionally or structurally important protein regions, such as ligand interaction interfaces or protein core. Additionally, we apply RINminer to several large and diverse protein families – eukaryotic proteases, extended AAA-ATPase domain, as well as viral RdRPs and jelly-roll capsids – and discover known as well as novel structural motifs.

### 1.1 Motif discovery approaches

In the past two decades, a variety of approaches have been developed to tackle the problem of structural motif discovery that are summarized below. First, we focus on methods that operate on structural information in a more direct fashion, and later we outline methods that, like our approach, use a graph-based abstraction of 3D structures.

One of the first methods to include structural information was SPratt [6, 7]. It combines the sequence motif discovery tool Pratt [8] with checks for structural conservation. Motifs here are represented as structurally conserved residues in sequence order. This introduces the limitation that motifs have to follow the same sequence order in different structures. Trilogy [9] starts with enumerating all spatially conserved patterns of three residues and then combines these three-residue patterns into larger patterns until a certain significance threshold is reached. This approach however introduces the requirement that each residue included in the motif must have a conserved distance to at least two other residues in the motif. In PINTS [10], common residue patterns are enumerated by step-wise inclusion of the residues lying closer than a distance threshold so that the superimposed patterns do not exceed a root mean square deviation (RMSD) significance cutoff. TerMo [11] approaches the problem by first defining groups of residues that are all in contact with each other and then identifying regular spatial motifs built by such groups.

Other approaches concentrate on the identification of very small structurally conserved motifs. Johansson et al. [12] for example only consider 4-6 non-contiguous amino acid residues, that can be found with simple enumeration. In DRESPAT [13] the authors define structural patterns as groups of 3 to 6 residues all within a certain distance from each other (a 12 Å cutoff is used) and compare them based on pairwise C_*α*_-C_*α*_ and C_*β*_-C_*β*_ distances as well as the distances between functional atoms. Wang and Scott [14] define a structural pattern as small as one amino acid residue and its spatial surrounding within a certain distance and build a kernel that allows for protein function classification. There are also approaches that use very specific definitions of motifs. Rahat et al., for example, define a motif as six nodes representing residues connected by covalent bonds or backbone hydrogen bonds [15]. Such motif definitions however suffer from the lack of generality outside their intended purpose.

Some approaches define motifs based on itemsets (groups of items, in this case of amino acid types) that occur within a small region of the structure, without considering sequence order. In these types of motifs only the presence or absence of a type of amino acid is considered, not the number or order of occurrences. An example of this are spatially cohesive itemsets [16] which are itemsets of three amino acid types that are found within a sphere of a certain size in at least a given number of structures.

Another group of methods look for conserved arrangements of larger structural motifs, so-called super-secondary structures, that consist of several consecutive secondary structure elements. Chiang et al. [17], for example, explore mutual arrangement of *β*-strands in *β*-sandwich proteins and create a hierarchical classification of their structures on this basis. Fernandez-Fuentez et al. [18] define a minimal super-secondary structure consisting of two consecutive secondary structures elements and then use binned distances and angle between them to classify these motifs and look them up in 3D structures of other proteins.

Some methods use statistical learning to identify structural motifs that can discriminate well between protein functional families. For example, GASPS [19] uses a genetic algorithm that learns the best such discriminating motifs defined in terms of their C_*α*_ coordinates and coordinates of centers of masses of the residues. Finally, some tools include information not only about protein sequence and 3D structure, but also about protein modeled dynamics such as the method presented by Chen and Bahar which classifies serine and cysteine proteases into subfamilies [20].

### 1.2 Graph-based representations of protein three-dimensional structures

Approaches that, like our approach, use graph representations of the protein 3D structures are a subset of the methods for finding structural motifs. Using graphs instead of considering all pairwise relations between residues to represent protein structure provides a way of reducing the residue-to-residue distances or contacts one has to consider. It also allows for more flexibility of the structures. There are multiple ways of obtaining such graph representations. Typically vertices are used to represent the individual residues of the protein, which are labeled with the type of amino acid. Some methods however rely on a representation at the level of individual atoms [21] or entire secondary structure elements [22] instead.

Edges between two vertices indicate the structural relationships between the corresponding residues and can have a label detailing this relationship. In its simplest form this relationship can be based on proximity in the 3D structure with a fixed distance cutoff and discretized distances as labels. The drawback of this definition is creating many spurious edges that connect residues not really contacting each other in the corresponding protein 3D structure due to steric occlusion by other residues.

A more sophisticated approach using almost-Delaunay tessellations was introduced by Huan et al. [23]. This approach reduces the number of edges compared to the proximity approach by only considering edges based on Delaunay tessellation [24]. Here the C_*α*_ atoms of the residues are taken as points and an edge between them is added if there exists a sphere such that the two points lie on its surface and no other point is inside the sphere. An almost-Delaunay tessellation also allows for small deviations from this classical Delaunay tessellation by tolerating shifts of all involved points by a certain small margin in order to fulfill the Delaunay tessellation criteria. Allowing these deviations helps to account for small structural variations in the data set.

Another approach of defining edges are so-called *residue interaction networks (RINs)*. In these networks, edges are constructed based on physical interactions between atoms of the individual residues. Doncheva et al. [25] made use of the established probe [26] and reduce [27] tools to determine such interactions and generate RINs. This is the representation we have chosen for this work, because it faithfully represents physical and chemical foundations of protein 3D structures. For this reason we have also chosen to use exact distance values as edge labels which allows us to more accurately determine the structural conservation of a motif. This however also requires the algorithm to allow approximate matching of edges.

### 1.3 Graph mining-based approaches

There exist a variety of approaches to identify biologically important motifs from such graph-based structure representations. Here we will focus on methods, that like our approach, are based on frequent subgraph mining (FSM) and related approaches. FSM, which will be explained in more details in Methods and Materials, is the enumeration of all subgraphs that occur in at least a certain number of graphs in a collection of graphs. In the case of structural motifs such a collection would correspond to a family of proteins, for example.

These methods often include modifications of the core frequent subgraph mining idea by introducing additional criteria to help better identify subgraphs representing potentially biologically relevant substructures. One previously mentioned study used discretized Euclidean distance labels on edges from almost Delaunay tesselation to include additional structural information in the graphs [23]. The idea behind such an approach is that functionally important motifs correspond to more strongly conserved geometric patterns, since their three-dimensional arrangement is essential for the protein function. Hence, if the corresponding edges in the graphs are labeled with Euclidean distances, subgraphs related to the protein function are expected to correspond to more similar edge labels across different structure graphs than non-relevant frequent subgraphs. Therefore requiring similar distance labels for the edges increases the specificity towards biologically meaningful subgraphs. Later this method was further improved by using multiple edge labels from overlapping discretization bins with edges being considered matching if there is an overlap of labels to account for similar distances on different sides of a discretization border [28]. This approach is similar in spirit to the method presented in this paper, but with the downside of not differentiating between bigger and smaller distance differences, if the distances fall into the same bin. Other studies have attempted to account for missense mutations, i.e. differences on the vertex labels, either by using post-processing, such as combining topologically identical subgraphs with differing vertex labels based on substitution matrices [29] or by including substitution probabilities in the frequent subgraph mining algorithm itself [30].

Finally, there is also a group of approaches that uses statistical properties of the subgraphs and is less specific to protein structures. Coherent subgraph mining [31] for example introduces the use of the mutual information between a subgraph and its subgraphs to identify meaningful subgraphs. Another approach is to mine for significant subgraphs [32], that is to only consider subgraphs that are significantly discriminative as a binary classifier based on the presence or absence of a subgraph. It can be reformulated as a problem of structural motif discovery by selecting the two classes to be within and outside a protein family. The idea has also been extended to select the set of subgraphs that in combination provide the best classification [33].

## 2 Methods and Materials

### 2.1 Graph terminology definitions

In this work we use a graph-based algorithm to detect conserved patterns in protein 3D structures. This algorithm is based on the concept of subgraph isomorphism. We define a graph *S* as a subgraph of graph *G* if *V*(*S*) ⊆ *V*(G) and *E*(*S*) ⊆ *E*(*G*). A graph *H* is isomorphic to a graph *G* if |*V*(*H*)| = |*V*(*G*)|, |*E*(*H*)| = |*E*(*G*)| and there exists a bijective function *ϕ*: *V_H_* ! *V_G_* such that *λ*(*v*) = *λ*(*ϕ*(*v*)) for every vertex *v* in *V*(*H*) and both (*ϕ*(*v*_1_), *ϕ*(*v*_2_)) ∈ *E*(*G*) and *λ*(*v*_1_,*v*_2_) = *λ*(*ϕ*(*v*_1_), *ϕ*(*v*_2_)) for every edge (*v*_1_,*v*_2_) in *E*(*H*) hold true given the labels *λ*. Following this a graph S is considered subgraph isomorphic to graph G if there exists a subgraph H of G that S is isomorphic to. We call a graph connected if there exists a path from every vertex to every other vertex. A spanning tree of a graph G is an acyclic and connected subgraph that covers the entire vertex set of G. We only consider connected undirected graphs without self-loops or multi-edges.

### 2.2 Residue interaction networks

Residue interaction networks are a way of describing protein structures as graphs of residues interacting or in contact with each other. The RINs we use in this project are constructed by representing each residue of the structure as a vertex labeled with its amino acid type. Edges are added between vertices that correspond to residues consecutive in sequence and are labeled as sequence-based interactions. After this, edges labeled as contact-based interactions are added for interactions determined by RINerator [25] if there is not already a sequence interaction between these residues. RINerator identifies interactions based on van der Waals surface contacts calculated by probe [26]. Further, all edges are labeled with the distance between the C_*α*_ of the corresponding residues in the structure. This means each edge has a two-dimensional label with the first element (*λ*_*type*_) representing the interaction type and the second (*λ_dist_*) the distance between the interacting residues.

### 2.3 Frequent subgraph mining

Frequent subgraph mining describes the process of determining frequently occurring subgraphs in a set of input graphs that are also referred to as the *graph database*. A subgraph is considered frequent or *supported* if its support value, i.e. the number of database graphs it is subgraph isomorphic to, is above a user-defined threshold. There exist multiple established algorithms to perform this task, such as gSpan [34], Gaston [35] and FFSM [36]. We base our approach on gSpan, which we extended with a number of new features in a custom implementation called RINminer.

The gSpan algorithm makes use of the concept of depth first search (DFS) codes to represent subgraphs. In the DFS code of a subgraph there is an entry for each edge of the graph with the two connected vertex IDs, followed by the two corresponding vertex labels and the edge label. The IDs of the vertices correspond to the traversal order of a spanning tree of the graph in DFS order. Edges that are part of the spanning tree are referred to as forward edges and go from a lower to a higher vertex ID. Similarly, edges that are not part of the spanning tree are referred to as backward edges and go from a higher vertex ID to a lower one. These backward edges occur when there are cycles in the graph.

gSpan also defines a total order on the entries in the DFS code based on whether the compared edges are forward or backward edges, the vertex IDs and the labels. This total order is then used to define a total order on the different DFS code representations which correspond to different spanning trees and traversal orders of a graph. For this only the first differing entries between two DFS codes are compared. The DFS code representation which is considered the smallest possible is the canonical DFS code for a given graph. By considering only subgraphs with canonical DFS codes, gSpan avoids duplicates. An example graph together with its canonical DFS code representation is shown in Fig 1.

**Figure 1:**
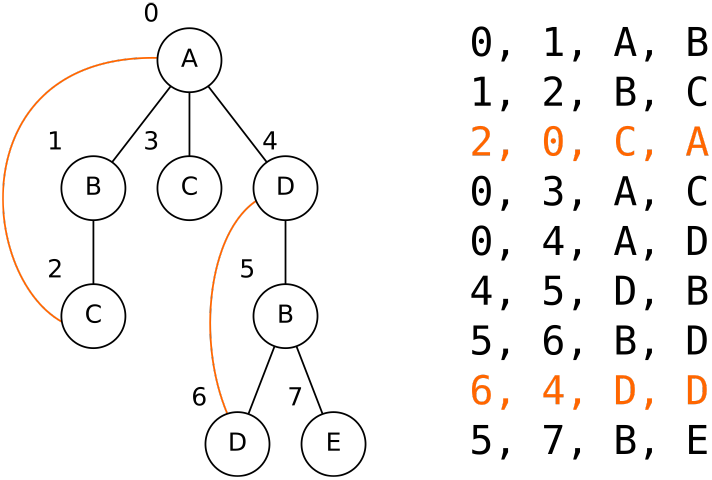
Example graph together with its canonicals DFS code representation. Shown is an example graph next to its canonical DFS code representation with backward edges highlighted in orange and vertex IDs shown next to each vertex. Edge labels are omitted.

After initialization, gSpan uses a supported single edge graph as starting point and proceeds with step-wise extension by either attaching a single edge-vertex pair (forward edges) or by connecting existing vertices in the graph (backward edges). All possible extensions considered in this step are determined by mapping the subgraph back into each database graph and detecting missing edges to vertices adjacent to vertices on the rightmost path in the DFS traversal. This extension process is repeated until the support of the extended subgraph falls below the specified threshold or its DFS code stops being canonical, at which point the algorithm backtracks to the last supported subgraph for which not all extensions have been explored. Once all supported extensions of the initial single edge graph have been explored, gSpan repeats this for the remaining supported single edge graphs. To determine whether a generated subgraph representation is canonical, the DFS code is compared to the smallest possible DFS code of this subgraph. The smallest possible DFS code of a subgraph is generated by starting with an empty graph and choosing the smallest possible extension leading to this subgraph at every step.

In RINminer, we parallelized this traversal of the search space with a level-wise partitioning scheme using subgraph pools. A pool contains all supported subgraphs of a certain edge-size. All subgraphs of edge-size *n* + 1 can be generated from the pool of subgraphs with edge-size n in parallel. The initial pool of subgraphs is generated by determining all single edge subgraphs from all database graphs. This parallelization scheme is possible because of the approach taken by the gSpan algorithm to detect and avoid duplicate subgraphs. It defines a canonical representation of subgraphs which depends on the order of construction of the subgraph. Only subgraphs constructed in the canonical order are considered, so the algorithm does not require comparison with other generated subgraphs. An example iteration in this parallelization scheme with our additions to gSpan is shown in Fig 2. Buehrer and Parthasarathy [37] suggest that adaptive partitioning is superior at balancing the processor load, however the level-wise approach introduced here proved to be beneficial for our score comparison scheme which we will describe later.

**Figure 2:**
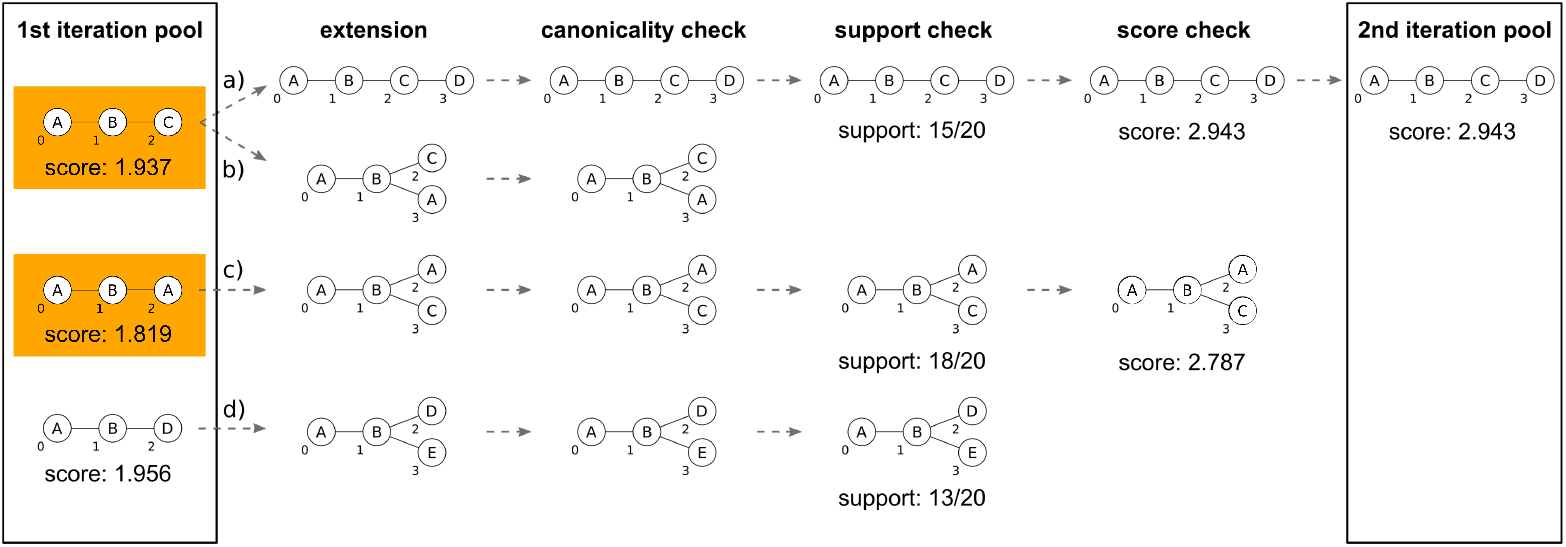
Outline of an example RINminer iteration. Shown is an example of the second iteration of RINminer which uses the pool of subgraphs generated during the first iteration to generate a new pool of subgraphs. First all possible extensions are determined followed by the different checks applied by RINminer assuming a support threshold of 15/20 and a minimum score increase of 0.9. Vertex IDs are shown to the bottom left of each vertex. Edge labels and subgraph variants that only differ in their edge labels are omitted for legibility. Extension *a* results in a valid subgraph which is added to the pool for the next iteration. Extension *b* is discarded due to a non-canonical representation. The canonical representation of the same subgraph is found in *c*. Extension *c* is discarded due to the score difference to its first parent subgraph. This subgraph’s parent subgraphs are marked with an orange background in the pool. Extension *d* shows a subgraph discarded due to low support.

To improve the performance of calculating the support of a generated subgraph in the extension phase we employ a hash table by calculating a hash of the DFS code representation of the extension. When determining all allowed extensions for a given subgraph and a given database graph this allows fast lookups of previous occurrences of the same extension found in other database graphs. This method can however only be used for exactly matching labels.

As an additional performance improvement over the original gSpan implementation, we store subgraph mappings as embedding lists similar to the approaches outlined by FFSM [36] and Gaston [35]. Embedding lists provide a space-efficient way to use subgraph mappings determined for a subgraphs of size *n* — 1 as starting points for mappings of a subgraphs of size *n* and share the memory for common parts.

New features of the algorithm added in RINminer include support for multidimensional edge labels, which allows us to consider both the residue distances as well as the interaction types in our RINs. Importantly, we allow for small deviations of the distance-based edge labels by introducing a tolerance when mapping subgraphs to database graphs. In addition to that, we introduce a scoring function based on these deviations to find the subgraphs corresponding to the structurally most conserved residue interactions. This scoring function is also used to limit the search space. We will now describe the two additions in more detail.

### 2.4 Approximate matching of distance labels

Instead of using discretized distance labels like many of the previously mentioned approaches, we propose to use exact values and employ a certain tolerance when matching. Without such a tolerance, labels would rarely match exactly due to natural fluctuations of protein 3D folds even in conserved substructures resulting in an artificially reduced support of the corresponding subgraph. The advantage of our approach is that it prevents accidentally missing matches between two close values that fall into different bins of an arbitrarily chosen discretization intervals. Another solution [28] to this problem that has previously been proposed is to consider multiple discretized distance labels per edge and require only one label to match for edges to be considered matching. This however can lead to the opposite problem of edges being matched despite having a distance difference that is larger than the distance between two discretization boundaries, which translates to only weak structural conservation. We propose a conceptually different approach that avoid this shortcoming as well. In our approach we consider the edges (*x,y*) ∈ *E*(*G*_1_) and (*ϕ*(*x*), *ϕ*(*y*)) ∈ *E*(*G*_2_) matching with regard to subgraph isomorphism if *λ_type_*(*x, y*) = *λ_type_*(*ϕ*(*x*), *ϕ*(*y*)) and *d*(*x, y, λ, ϕ*) ≤ *θ* with *d* being the distance difference:

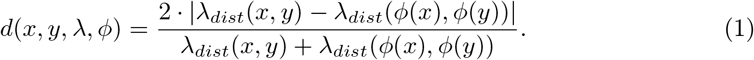

This definition is based on a related definition for distance similarity used by DALI [38] in its elastic similarity score to allow for higher variance in larger distances. Following DALI, we set the threshold *θ* to 0.2.

We apply this threshold only when determining the support of a subgraph, but not in the extension step of gSpan. The latter still requires exact label matches. In the following we will distinguish these two different types of matches by referring to them as *approximate* or *exact*, respectively. By only considering exact matches for the extension step, we ensure that we will only ever generate subgraphs that are exact subgraphs of at least one of the database graphs rather than combining edges from different database graphs. This still increases the total number of generated subgraphs by a factor proportional to the number of graphs in the database compared to a binning approach, but it avoids a potentially exponential growth that would occur when using approximate matching for the extension step. The impact of this is limited as the number of database graphs tends to be small (typically between 5 and 50) and also because we reduce the number of generated subgraphs with our scoring approach.

### 2.5 Subgraph scoring

The large number of subgraphs generated by FSM requires additional processing to determine the most interesting ones. Using the previously-described deviations in the distance labels we define a scoring function to determine the largest subgraphs that correspond to structurally strongly conserved residues. The score of a subgraph *S* for a database graph G given the sets of mappings Φ_*G*_ and the labels λ is defined as:

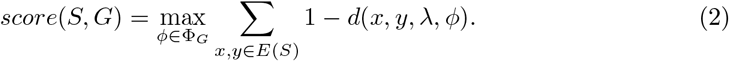

We then select the scores from the *t* database graphs with the highest scoring mappings, i.e. the t database graphs with the smallest total distance deviation for this subgraph and calculate the average as final score of the subgraph, where *t* is the gSpan support threshold. This limitation to t database graphs ensures comparability between subgraphs with different support values. Without it subgraphs that mostly have matches with high distance deviations would artificially improve their score if these database graphs were no longer matched after an extension. This property is important for our following use of the score which requires the ability to compare scores between two subgraphs with potentially different support values.

We make use of these scores to reduce the number of subgraphs generated by gSpan. In addition to the support requirement from the original gSpan algorithm, we also require the score to increase by a specified amount compared to all of its parent subgraphs. We used a value of 0.9 which was chosen such that it reduces the size of the search space while retaining the most relevant motif in the later discussed RdRP data set. As parent subgraphs we define all connected subgraphs of the subgraph that can be obtained by the removal of a single edge or a single edge-vertex pair, i.e. all subgraphs that could be extended to the subgraph in question ignoring DFS canonicality. Fig 2 shows the parent subgraphs of the example subgraph *c* indicated by an orange background. Using parent subgraphs makes this comparison independent from the order of construction, because all possible ways to construct this subgraph in a step-wise fashion without including disconnected graphs are considered. Further, we require all parent subgraphs to fulfill this requirement recursively. The subgraph pools introduced for parallelization allow us to look up these scores instead of recalculating them for every parent subgraph. It also allows for simple detection of parents not fulfilling the requirement, as such subgraphs would be missing from the pool. Using such an approach we only ever need to retain two such subgraph pools in memory at a time.

The parent subgraph lookup is implemented by determining removable edges or edge-vertex pairs according to the parent subgraph definition. Here we make use of multiple properties of DFS codes to implement this in an efficient manner. For this we first must relate our definition of edges removable due to cycles to the DFS code representation of a graph. This is given by the following theorem which is proven in S1 Proof:

#### Theorem 1.

In a graph with a given canonical DFS code representation the set of all edges cycles is the union of the sets of all backward edges together with the forward edges on the path connecting the end and starting vertices of each backward edge in the direction from the leaves to the root.

Using this theorem and the property of the canonical DFS code representation of a graph that all edges, backward or forward, originating from a vertex are sorted after the forward edge to that vertex, we can define an algorithm (Algorithm 1) to identify all removable edges in a graph a linear time. By iterating over the DFS code backwards we can count the number of cycles opened at a vertex by counting the number of backward edges originating from it. Similarly, we can count the number of cycles that will be closed once a vertex has been reached by the algorithm. In addition to the open cycles from backward edges originating from a vertex, open cycles are also inherited from child vertices in the spanning tree, i.e. forward edges originating from the vertex. We consider a vertex to be reached, once we hit the DFS code entry representing the forward edge pointing towards it. Using the number of open and closed cycles we can now for each forward edge determine whether there is currently an open cycle involving this edge. Backward edges are always considered removable, because they are known to be part of a cycle.

#### Algorithm 1 Determine all removable edges from a given graph and its canonical DFS code representation

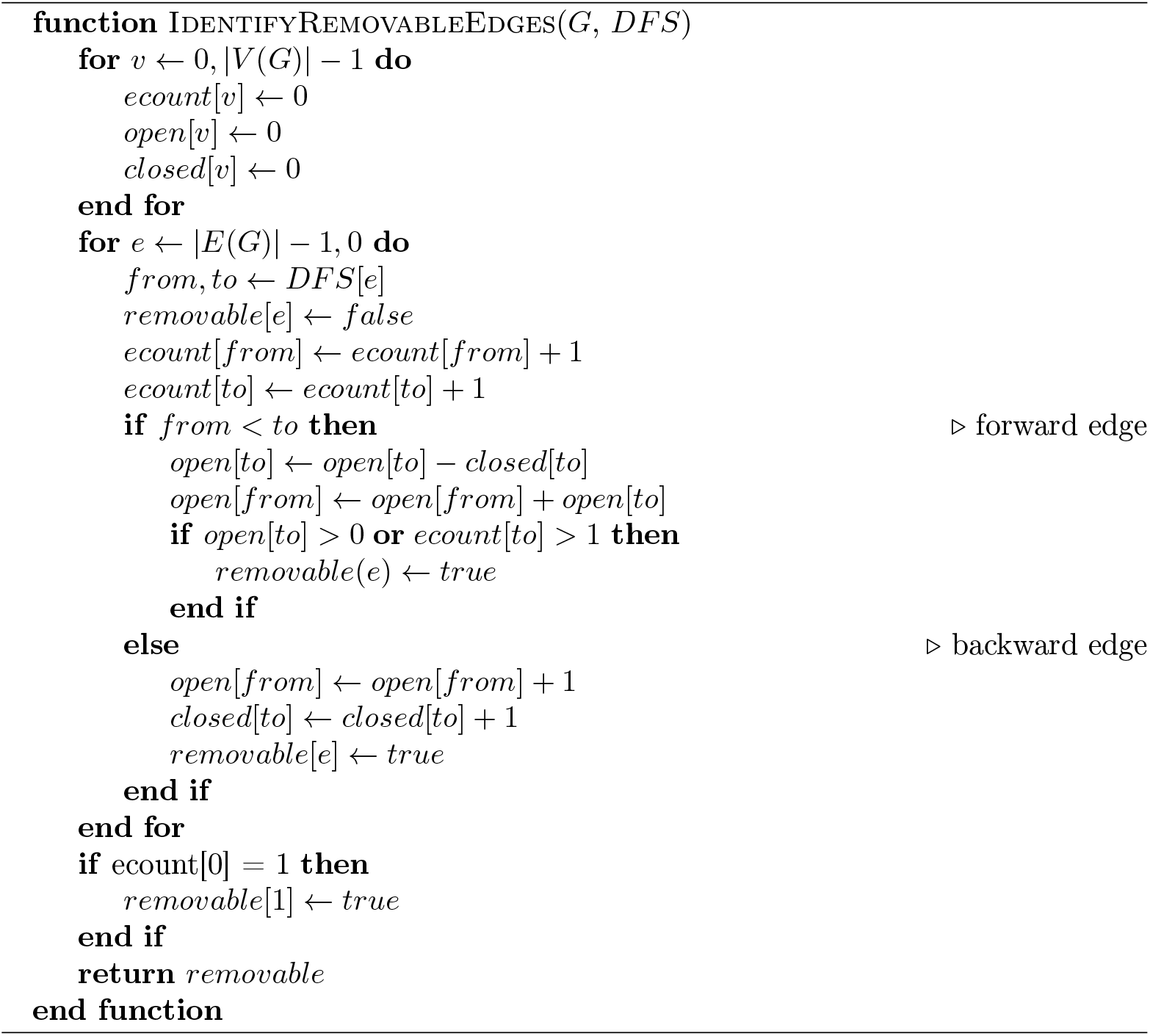

In addition to edges in a cycle, we also need to consider edges ending in terminal vertices. Here for both forward and backward edges, we just count the number of times outgoing and incoming vertices are encountered in the DFS code. For each forward edge we check if this count for the incoming vertex is 1, meaning that it is a terminal vertex and should be included as a removable edge-vertex pair. This works because in reverse canonical DFS code order the forward edge pointing towards a vertex is the last occurrence of that vertex with the exception of the root vertex. So as a special case in the first DFS entry (non-reverse order) the same is also done for the outgoing vertex, i.e. the root vertex.

With the list of removable DFS code entries corresponding to either edges or edge-vertex pairs, we now generate the parent graphs and determine their canonical DFS code representations. This is done analogously to the method used by gSpan to determine the canonicality of a DFS code representation by iteratively building the DFS code and choosing the edge resulting in the smallest DFS code entry. This DFS code representation can then be used for a hash-based lookup of the parent graphs in the subgraph pool.

### 2.6 PROSITE pattern data set and validation

We performed a validation of our approach based on protein families corresponding to patterns in the PROSITE database [4]. For this we determined the overlap between the residues matched by a pattern and the residues corresponding to subgraphs generated by RINminer for the same set of structures.

For each PROSITE pattern we selected all corresponding protein 3D structures marked as true positives and pruned the set of matching structures using PISCES [39]. We required the structures to be determined by X-ray crystallography at a resolution of at most 3.0 Å and we excluded structures that only contain C_*α*_ atoms to have reliable residue interaction information. The data set was then pruned to structure chains that share at most 50 % sequence identity. This identity threshold is higher than the distant relations our approach is intended for (≤ 30 %), which is due to the fact that PROSITE patterns represent sequence motifs found in more similar proteins. By using this less restrictive threshold we can avoid unnecessary further pruning of the already reduced data set and can later still filter out cases where the high sequence identity causes run time or memory issues due to too many subgraphs to explore. In a final step we prune data sets with less than three structures.

For each pattern we then ran RINminer with a support threshold ranging from min(3, 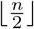) to *n*, where *n* is the number of structures remaining for each pattern after pruning. We then selected the 15 subgraphs with the highest subgraph score for each threshold. For each subgraph, only the highest scoring subgraph variant was used. A variant here refers to a subgraph with the same topology and vertex labels, but differing edge labels. We also excluded subgraphs and subgraph variants that were a part of already selected subgraphs with higher scores.

PROSITE patterns do not always specify exact sequences, but can also allow or disallow sets of amino acids for individual residues. Allowed amino acids in a pattern are encompassed by square brackets whereas disallowed are encompassed by curly brackets. Patterns can also contain wildcards, indicated by a lowercase x, which match every amino acid. To assess the overlap between a pattern and the subgraphs generated by RINminer, we use two different scoring functions considering the fact that subgraphs generated by RINminer cannot contain residues with multiple amino acid types. The first scoring function, which we refer to as the exact scoring function, only considers exact residue matches and ignores residues with multiple possible amino acid types. The second scoring function, on the other hand, includes residue matches against groups of amino acids and is in the following referred to as the non-exact scoring function. For both scoring functions the pattern is mapped onto the sequence given by the resolved residues of a structure. This may result in no mapping for structures with unresolved residues in which case these structures are ignored by the scoring function. It is also possible for a pattern to have multiple mappings. Next, the non-exact scoring function assigns a value of (20 — a)/19 to each matched residue in a mapping where *a* is the number of allowed amino acid types at a particular position of the pattern. In this way we include wildcard and amino acid group matches into the scoring function, but give them a lower contribution to the final score. For the exact scoring function, matched residues are assigned a value of 1 if *a* =1, and 0 otherwise. The maximum score of a pattern mapping in both scoring functions is defined as the sum of the values for all residues of the pattern. The score of a subgraph mapping is defined as the sum of the values of all residues mapped into the pattern normalized by the maximum score of the pattern mapping. In case of multiple subgraph or pattern mappings for a structure, the best score is used. The final score of a subgraph is the average of the best subgraph mapping scores for each structure.

For example the PROSITE pattern [IV]-{K}-[TACI]-Y-[RKH]-{E}-[LM]-L-[DE] (PS00226) has a maximum mapping score of 2 in the exact case and a score of 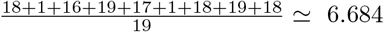 in the non-exact case. A subgraph mapping that only maps to the last two residues would have an exact score of 1 and a non-exact score of 1.947. The final scores for this subgraph assuming a single structure database would then be 0.5 and 0.291, respectively.

### 2.7 SCOP superfamily data set

Finally, we applied our approach to structures found in SCOP 1.75 [5] by generating a database for each superfamily. The pruning step and RINminer parameter calculation was done in the same way as for PROSITE but with a lower sequence identity threshold of 30 % when pruning to reflect the greater evolutionary distance found in superfamilies.

For this data set we also determined the significance of a subgraph using its classification power similar to Huan et al. [23]. Each subgraph was used to classify all structures from our pruned SCOP database. If a subgraph is approximately subgraph isomorphic to a database graph, this database graph and thus the corresponding protein structure was classified into the same superfamily as the originating database graph of the subgraph. The p-value of each subgraph was calculated using a onesided Fisher’s exact test on these classifications to determine the likelihood of a better classifier. When determining the significance of these p-values, we applied Bonferroni correction to account for multiple testing using a base threshold of 0.0001, following Huan et al. [23], and the number of data and parameter sets (3296) and the number of subgraphs evaluated for each of these sets (15), resulting in a significance threshold of 2.0 · 10^-9^.

### 2.8 Selected protein families data set

We also assembled a data set of three well researched protein families with known motifs and one family without any currently known structural motifs to evaluate the found motifs of our approach in more detail for these specific cases. The chosen families are AAA-ATPases, eukaryotic proteases, viral RdRPs and viral capsids. For the first two we again used SCOP but this time instead of taking entire superfamilies, we only focused on families (c.37.1.20 and b.47.1.2, respectively), but again a low sequence identity of at most 30 %. The last two data sets were adapted from data sets used by Caprari et al. [40] in their study of similar proteins in distantly related viruses. A list of chosen PDB structures can be found in Supplementary Table S2.

#### Implementation

RINminer was implemented in C. Scripts for generation of the data and analysis of the results were written in Python using Biopython [54] and SciPy [55]. The code is available at https://github.com/kalininalab/rinminer.

## 3 Results and Discussion

### 3.1 PROSITE Patterns

We generated 254 data sets consisting of between 3 and 7 RINs, where each data set corresponds to a single PROSITE pattern. After applying our method to these data sets, we excluded 126 data sets for which the enumeration of supported subgraphs had not terminated after 1 hour of runtime or run out of memory. The longer runtimes in comparison to most of our other experiments are caused by the higher similarity (50% identity cutoff) between the sequences, which then in turn leads to more similar RINs with a higher number of common subgraphs. For the remaining 128 data sets, we applied our scoring approach to evaluate the overlap between the known PROSITE patterns and the subgraphs identified by our approach. In each case we selected the largest frequent subgraph with the highest subgraph score (Equation 2).

When considering the 15 top-ranking subgraph across various support ranges (between half the size of the graph database and the size of the graph database), we observed that the ones with the highest subgraph score when comparing to the PROSITE patterns overlap with 48.7% (718 of 1475) of strictly conserved pattern residues, and the ones with the highest exact score cover 55.3% (815 of 1475) of them. For individual subgraphs with the highest subgraph score, their PROSITE exact scores have a bimodal distribution with 39.3% (48 of 122 patterns with at least one strictly conserved residue) with a value of 1, i.e. all strictly conserved residues included in the subgraph and 32.0% (39 of 122) with a value of 0, i.e. none of the strictly conserved residues included, and non-exact PROSITE scores have a median value of 0.39 (Fig 3A). Of the subgraphs with the highest PROSITE scores for each pattern across different support threshold values, 45.9% (56 of 122) have an exact score of 1 and 19.6% (24 of 122) of 0, and the median of non-exact scores is 0.49 (Fig 3B), indicating a high degree of overlap between the identified structural motifs and PROSITE patterns.

**Figure 3:**
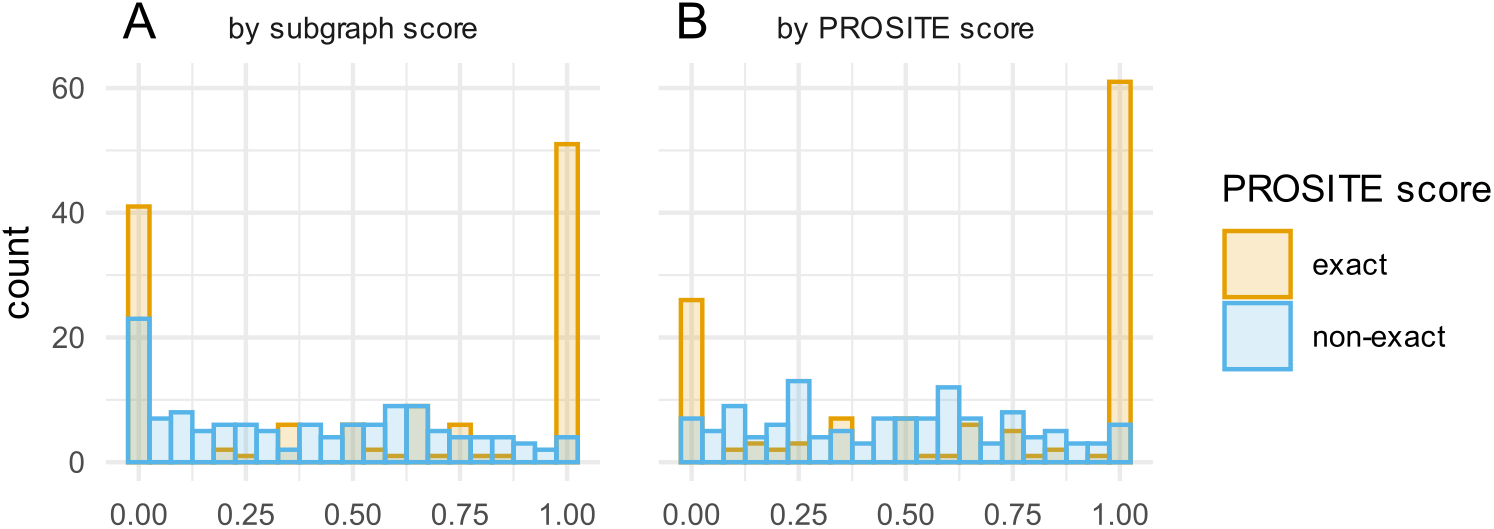
PROSITE score distributions for the best ranking subgraphs of each PROSITE pattern. (A) Subgraphs ranked by their subgraph score. (B) Subgraphs ranked by the exact/non-exact PROSITE score.

An example of a successful prediction is the Zinc finger RanBP2-type signature pattern ZF_RANBP2_1, where the top-ranking subgraph coincides with the pattern,which, in turn, contains only conserved positions (Fig 4A). Structurally, they also perfectly align, despite a relatively low average sequence identity within this family (only the residues of the PROSITE pattern are conserved in the corresponding domain, which corresponds to 6 out of 30 residues). An example of a more flexible pattern is the uroporphyrin-III C-methyltransferase signature SUMT_1, which contains strictly conserved, as well as variable positions (Fig 4B). Here our top-ranking subgraph includes three of four strictly conserved and three of eight variable pattern residues. Additionally, the subgraph includes four residues from a different pattern SUMT_2, which occurs in the same proteins, but is distant in sequence from SUMT_1. Here we demonstrate that two disjoint in sequence functional patterns come close to each other in space and can be included in a single structural motif.

**Figure 4:**
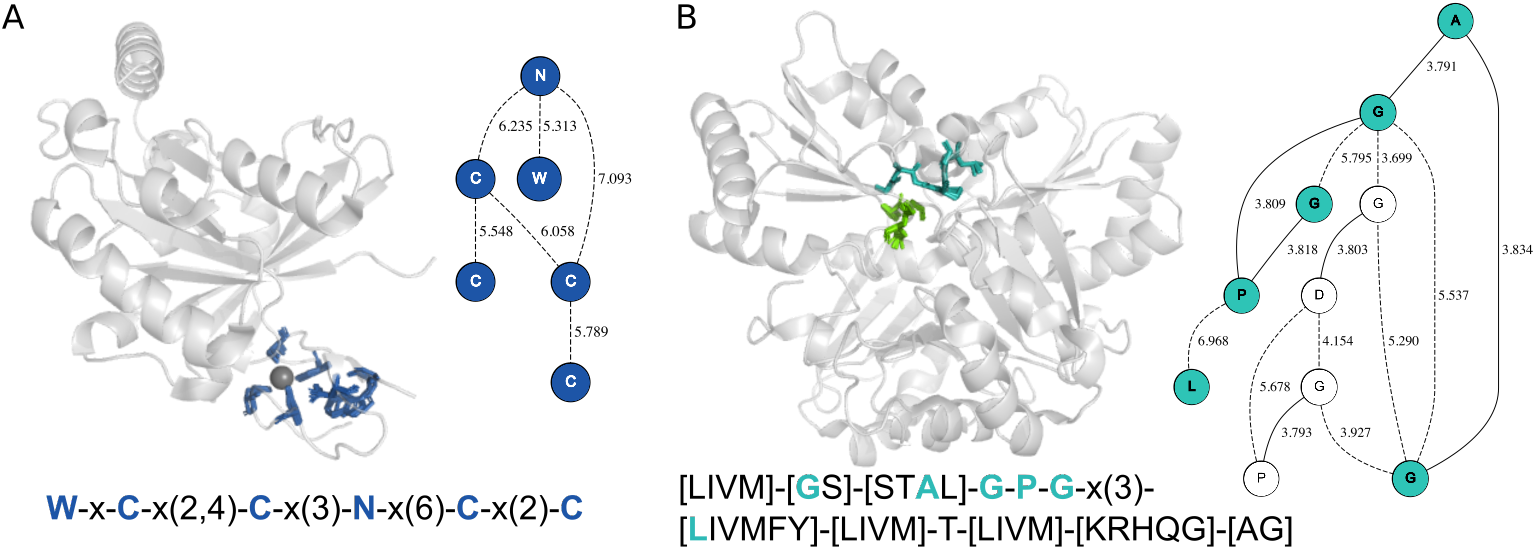
Examples of identified subgraphs for PROSITE pattern and their mapping into representative 3D structures. (A) PROSITE pattern ZF_RANPB_2 representing zinc finger RANBP2-type motif and its mapping into the structure of RanGDP-Nup153ZnF2 complex from human (PDB id 3GJ3). The overlapping residues between the motif and the subgraph are shown in dark blue. The Zn atom is shown as a gray sphere. (B) PROSITE pattern SUMT_1 representing the uroporphyrin-III C-methyltransferase signature 1 and its mapping into the structure of Precorrin-4 C11-methyltransferase from *Rhodobacter capsulatus* (PDB id 3NDC). The overlapping residues between the motif and the subgraph are shown in teal.

### 3.2 SCOP superfamilies

We applied our approach to 537 SCOP superfamilies, where we could find at least three representative structures of proteins with pairwise sequence identity up to 30% and a length of at least 50 residues using a filtering approach based on the PISCES algorithm [39]. With the chosen ranges of support thresholds this resulted in 3296 parameter sets, of which 120 did not complete within a specified time frame of 20 minutes or exceeded a memory usage of 1TB. To evaluate the found subgraphs we first determined how well the identified subgraphs can differentiate between structures of proteins belonging to the same and to different superfamilies. For that we implemented a classifier in a manner similar to Huan et al. [23] (see Methods and Materials for more details). Of the up to 15 top-ranked subgraphs in all 3296 parameter sets, 37.8% (9854/26068) are significantly classifying, i.e. the classification p-value is lower than the Bonferroni corrected significance threshold of 2.0 · 10^-9^. If we consider only the subgraphs with the highest score for each support threshold and each family, 41.9% are significantly classifying. Considering only the subgraph with the lowest p-value for each superfamily this yields 77.5% (396/511) significantly classifying subgraphs. These results indicate that our approach can identify subgraphs bearing some specific features that distinguish SCOP superfamilies from one another.

For SCOP, we do not have systematic description of the true functional motifs, although SCOP superfamilies were constructed in such a way that proteins in them share same functional mechanism and presumably common evolutionary origin [5]. We estimated the functional importance of the residues in identified subgraphs using a recently introduced structural annotation tool StructMAn [41]. Briefly, StructMAn explores structural environment of a given amino acid residue in a given protein, taking into account not only its own 3D structure, but also 3D structures of homologous proteins. Then it classifies residues into lying in the protein core, on the surface or on an interaction interface with another protein, low molecular weight ligand, DNA, or RNA. Residues in all these structural classes, except surface, are deemed to be functionally or structurally important. It has been previously shown that such a classification, despite its apparent simplicity, agrees well both with clinical annotations of mutations at individual protein positions and other computational tools for the prediction of functional consequences of mutations [41, 42].

We analyzed the distribution of the structural classification of all residues corresponding to significantly classifying top scoring subgraphs for different support thresholds (Table 3.2A). In order to be able to combine the results from superfamilies of different size, we consider the support threshold as a percentage of the numbers of structures in a superfamily. In these subgraphs, we observe a significant enrichment with residues classified as core, which may indicate their potential structural importance, and as interacting with ligands indicating their functional importance, whereas residues classified as surface residues are depleted (data not shown) across all support thresholds. A significant depletion can be observed in protein-protein interaction interfaces, which is a counter-intuitive result. Previously, it had already been observed that protein-protein interaction interfaces are not enriched in disease-associated mutation [42], and hypothesized that due to their large size, protein-protein interaction interfaces must possess a more complex functional landscape than being uniformly comprised of important residues. The percentage (0.6%-1.1%) of residues classified as DNA or RNA interacting rarely show a significant difference from the corresponding background percentage (1.2%) in the data set.

We also compared these results to the subset of residues that correspond to significantly classifying subgraphs that contain less than half typical frequent hydrophobic residues (A, I, L, V) and still have the highest score in the superfamily for a given support threshold (Table 3.2B). The removal of subgraphs consisting of mostly hydrophobic residues expectedly resulted in a depletion of residues classified as part of the core of the protein and an enrichment of residues interacting with ligands. This additional restriction therefore allows shifting the focus more towards functionally important motifs.

Another way of filtering the resulting subgraphs is to consider the rigidity of the corresponding motif. For this we also compared the classification distribution of the subgraphs for which the atoms from the corresponding residues in the different supporting structures can be superimposed with an average pairwise RMSD of at most 3Å in addition to the previous restrictions (Fig 5). Comparing the rigidity filtered results with and without the hydrophobic residue restriction (the last two plots in Fig 5A), shows that the hydrophobic residue restriction does not have a strong effect in this case as it does without the rigidity filtering (the first two plots in Fig 5A). This probably means that there are stricter geometric constraints on the position of functionally important residues in these sites than in protein core. When one considers that ligandbinding sites, defined as they are by StructMAn, can frequently be catalytic sites, one can hypothesize that identified patterns are related to catalytic residues.

**Figure 5:**
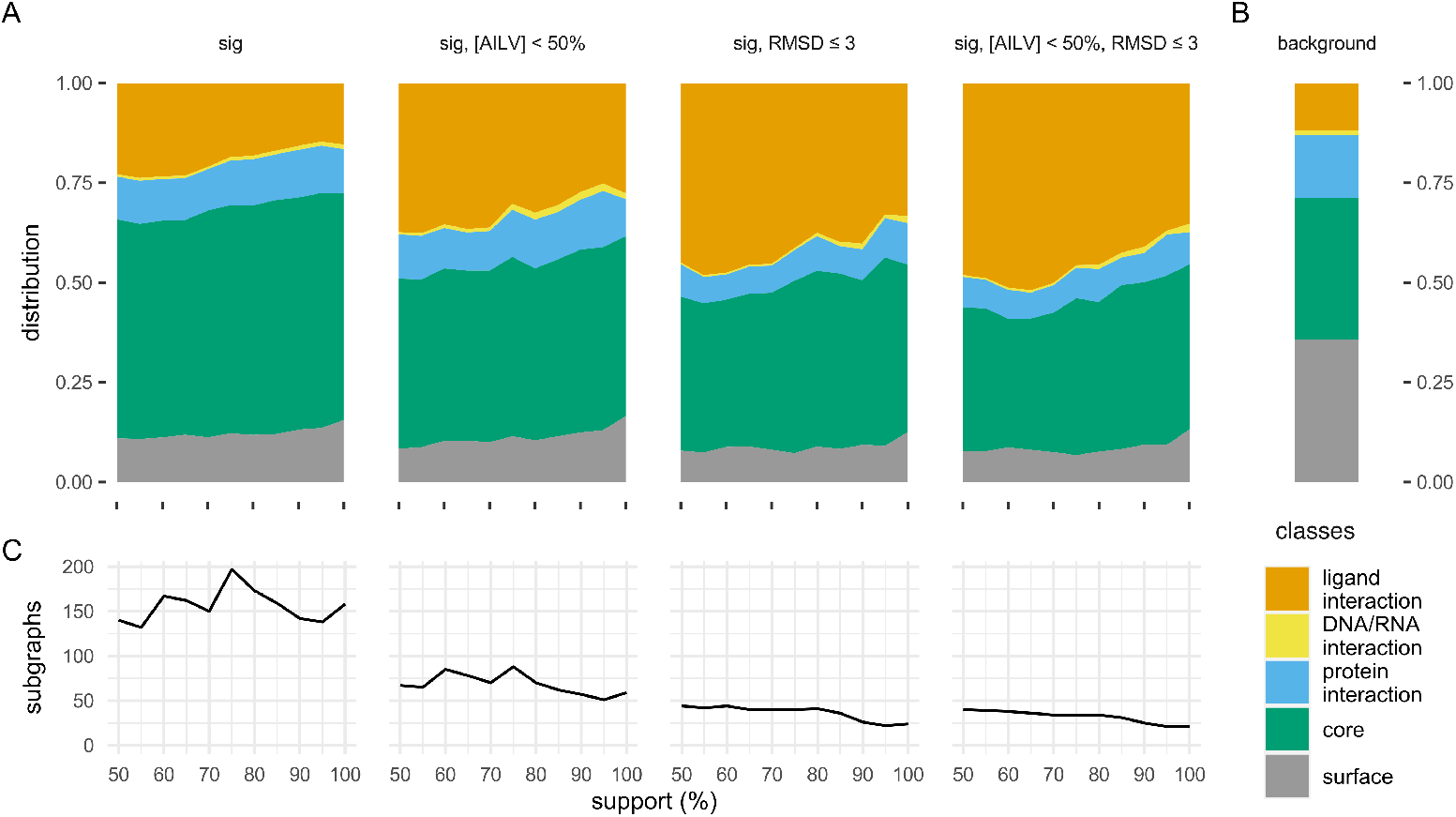
Structural classification of residues corresponding to identified subgraphs. (A) Stacked area charts of the distributions of structural classes for different categories of subgraphs at different support thresholds. (B) Background distribution across all residues in the data set. (C) Number of identified frequent subgraphs in each category of at different support thresholds. The support thresholds in A and B are in percent of the database size from 50% to 100% with a step size of 5% and the categories are from left to right: all significantly classifying subgraphs (sig); significantly classifying subgraphs that comprise less than a half hydrophobic residues A, I, L, V; significantly classifying subgraphs, whose mappings into protein 3D structures can be superimposed with a RMSD less than 3 Åon average (only the highest scoring mapping for each structure is considered); significantly classifying non-hydrophobic subgraphs with mappings that can be superimposed with a RMSD up to 3 Å on average.

In some cases the spatial location of the residues from a top-ranked subgraph can readily suggest their function. For example, for the SCOP superfamily e.3.1 (beta-lactamase/transpeptidase-like) we identify a subgraph that corresponds to a very well conserved structural motif (Fig 6A). This motif includes two lysines and two serines that can be separated by a great distance in the protein sequence (the number of residues between one of the lysines and the following serine can be between 44 and 245). The serine and lysine of the S-x-x-K submotif are parts of the beta-lactamase active site [43, 44]. In agreement with this, in one of the structures (PDB id 1XKZ), this motif defines a pocket where an antibiotic derivative acylated ceftazidime is bound. In other cases, e.g. for the rhodanese/cell cycle control phosphatase superfamily (c.46.1) we can identify a structurally very well conserved motif buried in the protein core (Fig 6B), which may play a role in building the protein scaffold.

**Figure 6:**
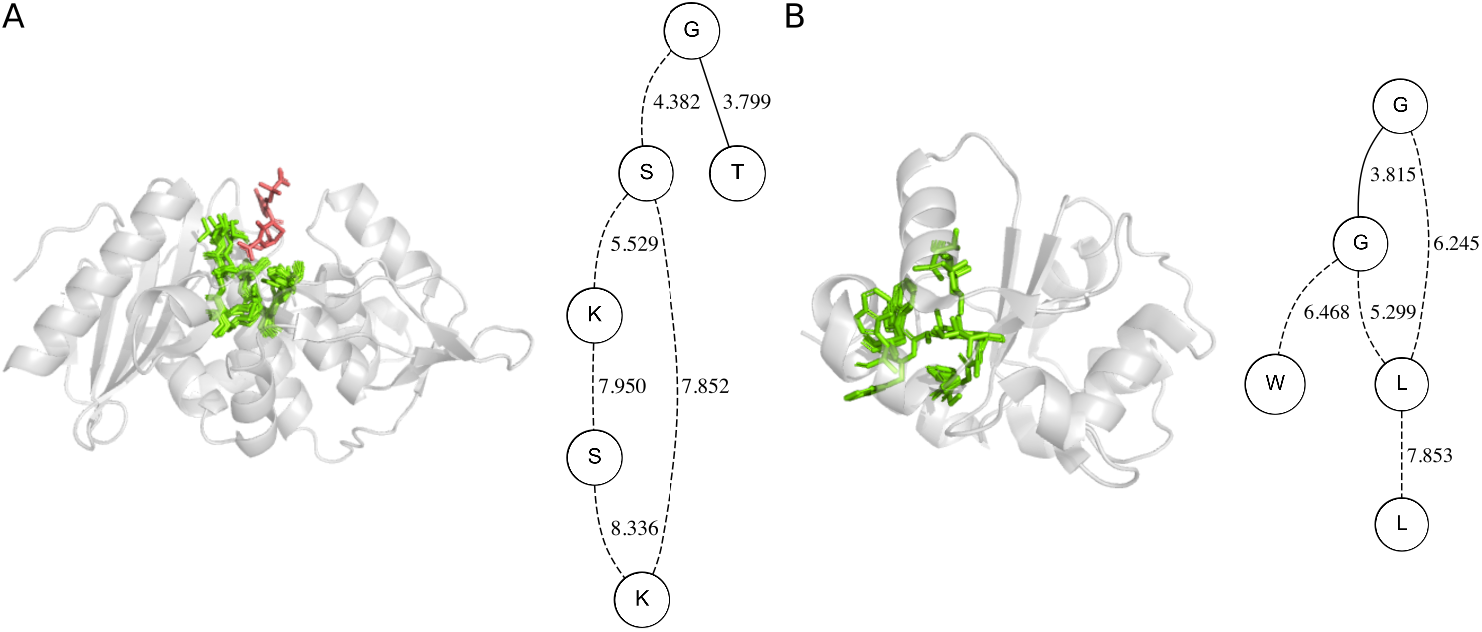
Examples of identified subgraphs for SCOP superfamilies and their mapping into representative 3D structures. Residues corresponding to the subgraph are shown as sticks and colored green. (A) SCOP superfamily e.3.1 (beta-lactamase/transpeptidase-like) and mapping into the structure of S216A mutant of *Streptomyces* K15 DD-transpeptidase (PDB id 1ES5). Inhibitor acylated ceftazidime from aligned structure of acylated beta-lactam sensor domain of Blar1 from *S. aureus* (PDB id 1XKZ) is shown in salmon. (B) SCOP superfamily c.46.1 (rhodanese/cell cycle control phosphatase) and mapping into the structure of a USP8-NRDP1 complex from human (PDB id 2GWF).

### 3.3 Selected diverse protein families

In addition to superfamilies, we also looked at two families in SCOP which show extremely diverse sequences (eukaryotic proteases b.47.1.2 and extended AAA-ATPase domain c.37.1.20). SCOP families are the next hierarchical level after superfamilies, usually proteins from the same family demonstrate an easily detectable sequence similarity. However, in the cases of eukaryotic proteases and extended AAA-ATPase domain they are extremely large and diverse, which makes them suitable for analysis with RINminer. Further, we analyzed two sets of structures from virus protein families which are of special interest due to their particularly diverse sequences. The first family comprises 3D structures from 12 manually selected RNA-dependent RNA polymerases (RdRP) from positive- and negative-sense single-strand, as well as from double-strand RNA viruses. The other family contains 49 3D structures of jelly-roll capsids from a very diverse set of virus families, including double-strand DNA (e.g. *Baculoviridae, Papillomaviridae, Polyomaviridae*), double-strand RNA (*Birnaviridae*), single-strand DNA (*Circoviridae, Parvoviridae*), and positive-sense single-strand RNA (e.g. *Bromoviridae, Caliciviridae*, several *Picornavirales* families) viruses. The average pairwise sequence identity in these four families ranges between 10.9% and 26.2% (Table 2).

**Table 1:** Distribution of the structural classification of the residues corresponding to the top scoring mappings of the top-scoring subgraphs for each superfamily at a given support threshold for the SCOP data set. Only considering significantly classifying subgraphs (A), only significantly classifying and containing less than 50% frequent hydrophobic residues (B), all residues in the data set (C). Shown for each structural class are the number of residues in the class (#) and percentage as well as the odds-ratio and significance (*** < 2.2 x 10^-16^, ** < 0.01, * < 0.05) according to Fisher’s exact test.

|  | sup | ligand interaction |  |  |  | DNA/RNA interaction |  |  |  | protein interaction |  |  |  | core |  |  |  |
| --- | --- | --- | --- | --- | --- | --- | --- | --- | --- | --- | --- | --- | --- | --- | --- | --- | --- |
|  |  | # | % | odds |  | # | % | odds |  | # | % | odds |  | # | % | odds |  |
| A | 50 | 1934 | 22.8 | 2.225 | *** | 54 | 0.6 | 0.521 | ** | 903 | 10.7 | 0.637 | *** | 4648 | 54.9 | 2.212 | *** |
|  | 60 | 2276 | 23.4 | 2.296 | *** | 62 | 0.6 | 0.520 | ** | 1018 | 10.5 | 0.622 | *** | 5297 | 54.4 | 2.170 | *** |
|  | 70 | 1977 | 21.0 | 1.997 | *** | 55 | 0.6 | 0.477 | ** | 979 | 10.4 | 0.619 | *** | 5351 | 56.8 | 2.399 | *** |
|  | 80 | 1648 | 18.2 | 1.664 | *** | 86 | 0.9 | 0.779 | * | 1046 | 11.5 | 0.695 | *** | 5220 | 57.5 | 2.468 | *** |
|  | 90 | 1068 | 15.7 | 1.394 | *** | 71 | 1.0 | 0.860 |  | 809 | 11.9 | 0.722 | *** | 3955 | 58.2 | 2.531 | *** |
|  | 100 | 1023 | 15.4 | 1.362 | *** | 76 | 1.1 | 0.945 |  | 729 | 11.0 | 0.659 | *** | 3775 | 56.9 | 2.398 | *** |
| B | 50 | 1388 | 37.4 | 4.485 | *** | 24 | 0.6 | 0.530 | ** | 410 | 11.0 | 0.663 | *** | 1581 | 42.6 | 1.341 | *** |
|  | 60 | 1673 | 35.4 | 4.135 | *** | 41 | 0.9 | 0.714 | * | 478 | 10.1 | 0.602 | *** | 2041 | 43.2 | 1.379 | *** |
|  | 70 | 1456 | 36.1 | 4.258 | *** | 38 | 0.9 | 0.776 |  | 399 | 9.9 | 0.587 | *** | 1731 | 43.0 | 1.364 | *** |
|  | 80 | 1045 | 32.5 | 3.608 | *** | 56 | 1.7 | 1.446 | ** | 394 | 12.2 | 0.746 | ** | 1385 | 43.0 | 1.367 | *** |
|  | 90 | 659 | 27.3 | 2.810 | *** | 47 | 1.9 | 1.622 | ** | 302 | 12.5 | 0.765 | ** | 1105 | 45.8 | 1.528 | *** |
|  | 100 | 655 | 27.6 | 2.856 | *** | 34 | 1.4 | 1.187 |  | 221 | 9.3 | 0.549 | *** | 1070 | 45.1 | 1.488 | *** |
| C | - | 136 105 | 11.8 | - | - | 13 946 | 1.2 | - | - | 181 324 | 15.7 | - | - | 410 091 | 35.6 | - | - |

**Table 2:** Average pairwise sequence identities of the selected protein families.

| Family | Number of structures | Average sequence identity |
| --- | --- | --- |
| Eukaryotic proteases b.47.1.2 | 7 | 26.2% |
| Extended AAA-ATPase domain c.37.1.20 | 23 | 14.6% |
| RNA-dependent RNA polymerases | 12 | 11.2% |
| Jelly-roll capsids | 49 | 10.9% |

The eukaryotic proteases are ubiquitous proteins with diverged sequences and 3D structures resolved for very many species. In our data set, we collected 7 3D structures of proteins with average sequences identity of 26.2%. Using our approach, we can identify a conserved structural motif that includes the catalytic triad [45] in 6 of them (Fig 7A). A somewhat larger common subgraph in this case also reflects the higher sequence similarity within this family. One structure (1A7S) from our data set not containing this motif is due to it having an inactivating H41S mutation [46].

**Figure 7:**
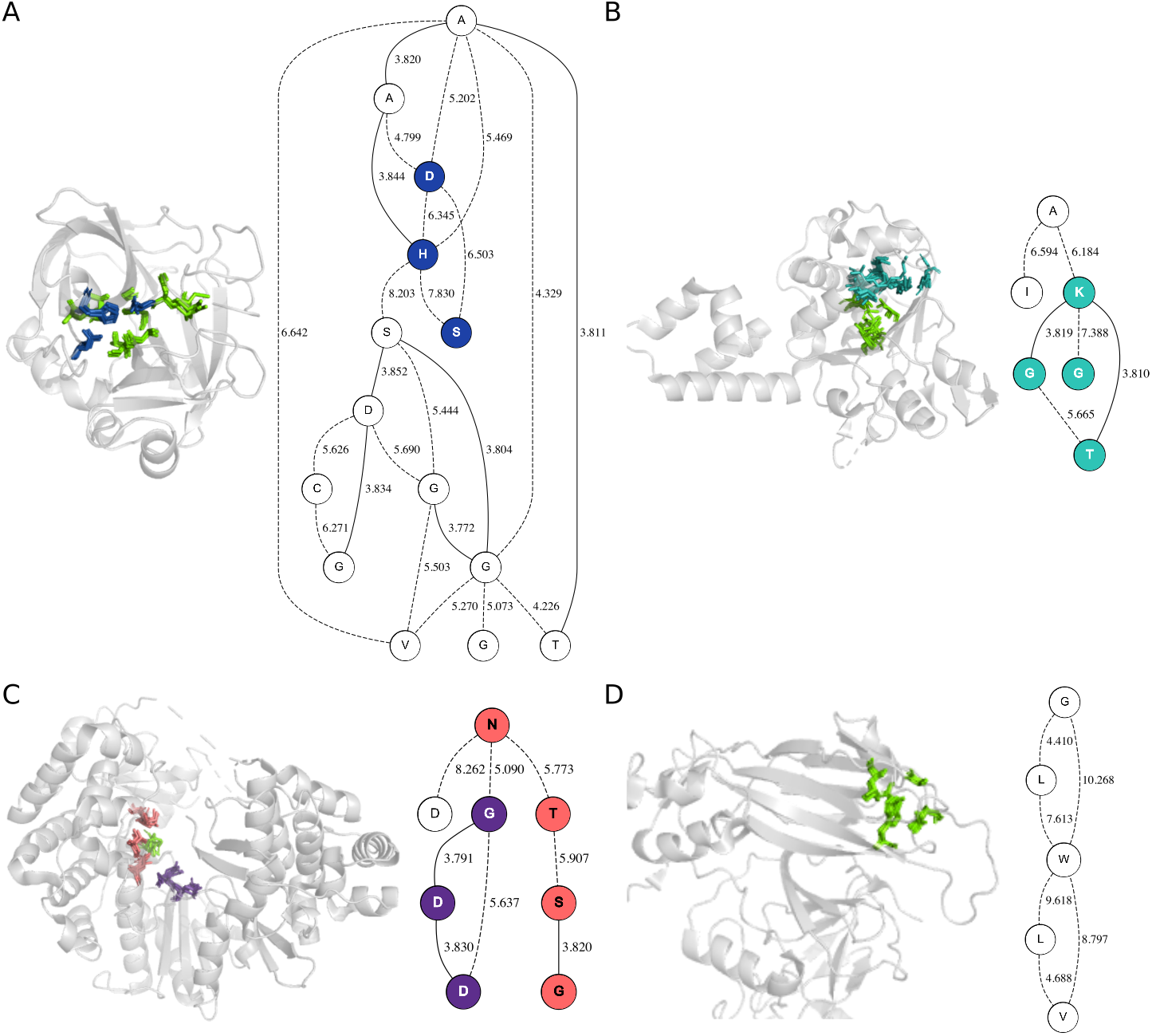
Examples of identified frequent subgraphs in selected diverse protein families and their mapping into protein 3D structures. (A) Eukaryotic proteases b.47.1.2. The catalytic triad is shown in dark blue. (B) Extended AAA-ATPase domain c.37.1.20. Walker A motif is shown in teal. (C) RNA-dependent RNA polymerases. Motif B is peach, motif C in purple. (D) Jelly-roll capsids.

The extended AAA-ATPase family is a large and diverse protein family involved in different cellular processes. In our data set, we selected 23 3D structures of AAA-ATPases with on average 14.6% sequence identity. Using our method, for the support threshold 11 we identify a subgraph corresponding to a known functional Walker A motif [47] involved in phosphate binding (Fig 7B).

Functional motifs in the RdRP family have been previously identified by sophisticated sequence comparison. One distinguishes seven functional motifs with relatively degenerate consensus sequences, which play different roles in catalysis and coordination of the substrate and lie at large distances that can vary substantially among viruses [2]. The subgraph with the highest score identified using the support threshold 6 defines a structural pattern that contains three of these motifs packed tightly together in the 3D space (Fig 7A): one of the two aspartate residues of the motif A that coordinate the metal ions required for catalysis [48], motif B, which is located in the active center of the polymerase and is responsible for distinguishing between ribose- and deoxyribose-based nucleotides [49], and motif C, which is a part of the active site and participates in coordination of metal ions as well [50, 51]. These motifs come from distant regions of the sequences, but form a single compact structural arrangement.

Jelly-roll capsids are found in very distant viral families, nevertheless their common evolutionary origin has been suggested, based on an exceptional conservation of their 3D structure [52]. There is very little sequence similarity among these structures: structure-based pairwise alignments of our 49 structures yield on average 10.9% sequence identity. For most support thresholds, we can identify only very small (up to 3 vertices) frequent subgraphs that consist of most common amino acids, and hence are perhaps not very interesting. However, for the threshold of 6 (12.2%), we find a subgraph of 5 vertices comprised of a tryptophan along with multiple hydrophobic amino acids (Fig 7B). Interestingly, in the 3D structure of the capsid, these residues form a very conserved pattern of interactions in the hydrophobic core, being perhaps a part of the proteins’ conserved structural scaffold as this motif shows a remarkably low average RMSD (0.354 Å). The 6 viruses we found this motif in belong to 3 distantly related families: *Senecavirus* and *Hepatovirus A* to *Picornaviridae, Cripavirus* and *Triatovirus* to *Dicistroviridae*, and *Vesivirus* and *Lagovirus* to *Caliciviridae*. These families have been assigned to the evolutionary related *Picornavirus-like superfamily* [53].

## 4 Conclusions

We propose a novel data-driven approach and the corresponding tool, RINminer, for the identification of functionally important motifs in distantly related proteins with known 3D structures. The basis of our approach is the technique of frequent subgraph mining that has previously been applied in life sciences, for example to identify common chemical moieties [34, 35] or protein structural motifs [23, 28].

A significant novel feature of our approach that has been implemented in RINminer for the first time is that we propose a concept of approximate label matching, in that subgraphs of graphs representing protein 3D structures are matched, even if the edge labels representing the Euclidean distance between proteins residues do not match exactly. We introduce an additional tunable parameter that governs the allowed deviation between the edge labels. This allows us to use exact Euclidean distances between protein residues directly, without binning that was employed in previous studies [28].

Another advantage is that we convert protein 3D structures to graphs using residue-interaction networks [25] that respect the physicochemical nature of interactions among protein atoms. Additionally, we differentiate between covalent and non-covalent interactions by labeling edges as *sequence* or *structural*. Both distance and interaction type are encoded as a multidimensional edge label. In the process of frequent subgraph mining only the dimension of the labels that represent the interaction type are matched exactly. This has been made possible by implementing support for multidimensional labels, which is also a unique feature of RINminer.

Due to run time constraints, only approximate matching of edge labels but not vertex labels has been considered, although approximate label matching for vertices is also supported by the tool. Using this option will, for example, enable mining for subgraphs that include not only exactly same, but chemically similar residues. Another way to account for physicochemical similarities between residues would be to use pseudoresidue labels, i.e. group several related residues together under one label (e.g. all aliphatic or all positively charged). We explored this possibility, but the resulting subgraphs were either too unspecific, or the runtimes were too long. Introducing approximate isomorphism on the residue labels would be thus a more flexible way to account for such relationships, which will be a matter of future research.

In various validation and application scenarios, we have shown that our approach is able to re-discover known motifs and propose novel ones. In particular, we suggested a novel structural motif in jelly-roll capsid proteins from several viruses of the picornavirus-like superfamily.

## 5 Acknowledgments

The authors would like to thank Nadezhda Doncheva for her assistance with RINerator and Alexander Gress for adapting StructMAn to make some of the analysis shown in this paper possible. We are also grateful to Alexander Gress and Fawaz Dabbaghie for critical reading of the manuscript.

## S1 Proof. Proof of Theorem 1

*Proof.* It is trivial to see that a path of forward edges where the first and last vertex of the path are connected by a backward edge forms a cycle, however it still has to be proven that no further edges than those stated by the theorem can be part of any cycle. Assume that there is a forward edge *e* = (*e_root_, e_leaf_*) that is part of a cycle with the vertex *e_root_* being closer to the root and *e_leaf_* closer to the leaf such that e is not considered part of a cycle by the theorem, i.e. it does not lie on a path connecting the end of a backward edge with its start. We also need to consider a backward edge *b* that is involved in the formation of such a cycle, because the spanning tree consisting of forward edges alone contains no cycles. Given that *e* is supposed to be part of a cycle, there needs to be a path that connects *e_root_* to *e_leaf_* without traversing e. Because in the spanning tree the only path connecting *e_root_* to *e_leaf_* is e, *b_root_* needs to be on the path between *e_root_* and the root vertex in order for such an *e_leaf_* to *e_root_* path to exist. Since we require *e* to contradict our theorem, *e* must not be on the forward edge path between *b_leaf_* to *b_root_*, which means that *b_leaf_* has to be on a branch that does not contain e, i.e. on a branch that branched off of the path between *b_root_* and *e_root_*. Now in order to complete a cycle there has to be a path between this branch and *e_leaf_* without going over e. This would require another backward edge. However by construction there can only ever be backward edges between the vertices on the path from the leaf to the root node (i.e. the rightmost path at the time of construction), so any backward edge would end on either the branch or another vertex on the path between *e_root_* and the root node resulting in one of the previous problems. Therefore such an edge e can not exist and the theorem holds true.

**Supplementary Table S2.**
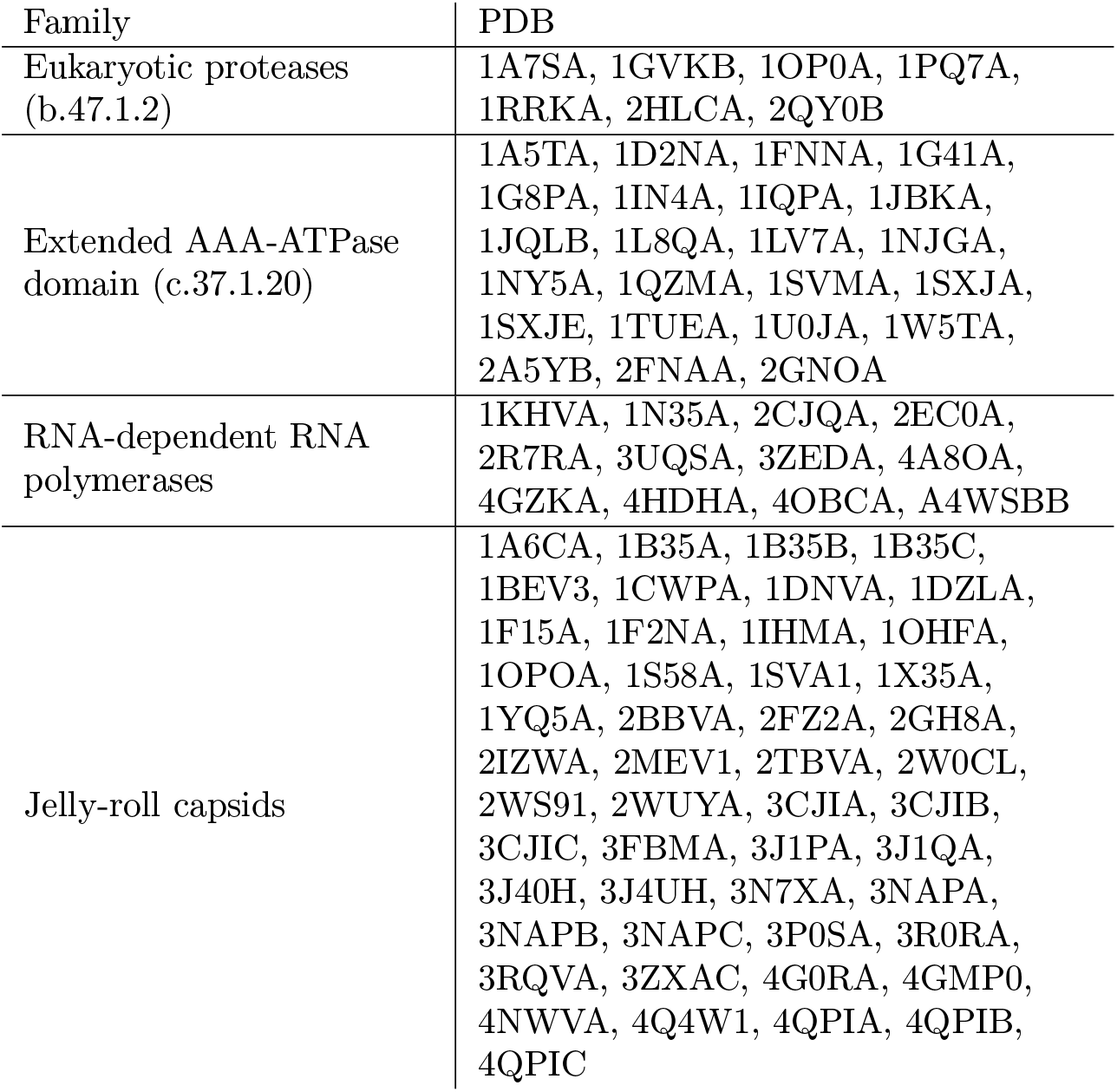
Structures used in the Selected protein families dataset. PDB identifiers are given followed by chain identifiers. Note that in some cases multiple chains were used for graph generation.

| Family | PDB |
| --- | --- |
| Eukaryotic proteases<br>(b.47.1.2) | 1A7SA, 1GVKB, 1OP0A, 1PQ7A,<br>1RRKA, 2HLC A, 2QY0B |
| Extended AAA-ATPase<br>domain (c.37.1.20) | 1A5TA, 1D2NA, 1FNNA, 1G41A,<br>1G8PA, 1IN4A, 1IQPA, 1JBKA,<br>1JQLB, 1L8QA, 1LV7A, 1NJGA,<br>1NY5A, 1QZMA, 1SVMA, 1SXJA,<br>1SXJE, 1TUEA, 1U0JA, 1W5TA,<br>2A5YB, 2FNAA, 2GNOA |
| RNA-dependent RNA<br>polymerases | 1KHVA, 1N35A, 2CJQA, 2EC0A,<br>2R7RA, 3UQSA, 3ZEDA, 4A8OA,<br>4GZKA, 4HDHA, 4OBCA, A4WSBB |
| Jelly-roll capsids | 1A6CA, 1B35A, 1B35B, 1B35C,<br>1BEV3, 1CWPA, 1DNVA, 1DZLA,<br>1F15A, 1F2NA, 1IHMA, 1OHFA,<br>1OPOA, 1S58A, 1SVA1, 1X35A,<br>1YQ5A, 2BBVA, 2FZ2A, 2GH8A,<br>2IZWA, 2MEV1, 2TBVA, 2W0CL,<br>2WS91, 2WUYA, 3CJIA, 3CJIB,<br>3CJIC, 3FBMA, 3J1PA, 3J1QA,<br>3J40H, 3J4UH, 3N7XA, 3NAPA,<br>3NAPB, 3NAPC, 3P0SA, 3R0RA,<br>3RQVA, 3ZXAC, 4G0RA, 4GMP0,<br>4NWVA, 4Q4W1, 4QPIA, 4QPIB,<br>4QPIC |

